# Genome and transcriptome analysis of the beet armyworm *Spodoptera exigua* reveals targets for pest control

**DOI:** 10.1101/2021.05.14.444148

**Authors:** Sabrina Simon, Thijmen Breeschoten, Hans J. Jansen, Ron P. Dirks, M. Eric Schranz, Vera I.D. Ros

## Abstract

**Background:** The genus *Spodoptera* (Lepidoptera: Noctuidae) includes some of the most infamous insect pests of cultivated plants including *Spodoptera frugiperda*, *Spodoptera litura* and *Spodoptera exigua*. To effectively develop targeted pest control strategies for diverse *Spodoptera* species, genomic resources are highly desired. To this aim, we provide the genome assembly and developmental transcriptome comprising all major life stages of *S. exigua*, the beet armyworm. *Spodoptera exigua* is a polyphagous herbivore that can feed from > 130 host plants including several economically important crops.

**Results:** The 419 Mb beet armyworm genome was sequenced from a female *S. exigua* pupa. Using a hybrid genome sequencing approach (Nanopore long read data and Illumina short read), a high-quality genome assembly was achieved (N50=1.1 Mb). An official gene set (OGS, 18,477 transcripts) was generated by automatic annotation and by using transcriptomic RNA-seq data sets of 18 *S. exigua* samples as supporting evidence. In-depth analyses of developmental stage-specific expression in combination with gene tree analyses of identified homologous genes across Lepidoptera genomes revealed potential *Spodoptera*-specific genes of interest such as mg7 and REPAT46 upregulated during 1^st^ and 3^rd^ instar larval stages for targeted pest-outbreak management.

**Conclusions:** The beet armyworm genome sequence and developmental transcriptome covering all major developmental stages provides critical insights into the biology of this devastating polyphagous insect pest species with a worldwide distribution. In addition, comparative genomic analyses across Lepidoptera significantly advance our knowledge to further control other invasive *Spodoptera* species and reveals potential lineage-specific target genes for pest control strategies.

## Background

Analysis of genome and transcriptome data can be used to study many important questions ranging from species-specific mutations to comparative genomic evolutionary patterns. The genus *Spodoptera* is known for the high number of notorious pest species causing enormous agricultural damage resulting in economic losses worldwide, including *Spodoptera exigua*, *Spodoptera frugiperda* and *Spodoptera litura* (Pogue 2002; Goergen, et al. 2016; Cheng, et al. 2017; EPPO 2017). The beet armyworm, *S. exigua* (Hübner) (Lepidoptera: Noctuidae) is a devastating polyphagous insect pest with a worldwide distribution (Mehrkhou, et al. 2012; Fu, et al. 2017), being able to feed on more than 130 plant species from at least 30 families including several economically important crops such as sugar beet, cotton, soybean, cabbage, maize and tomato (Merkx-Jacques, et al. 2008; Robinson, Ackery, et al. 2010; Mehrkhou, et al. 2012; Fu, et al. 2017). *Spodoptera exigua* originated in Southern Asia and was subsequently introduced to other parts of the world including North America and Europe (Mehrkhou, et al. 2012; Fu, et al. 2017). It is widely distributed in the tropical and subtropical regions and migrates into more temperate regions throughout the growing season (Pogue 2002). Its long-distance migration likely played a major role in the geographic expansion of populations and its spread across the world (Fu, et al. 2017). In temperate regions it can be abundant in greenhouses (Smits, et al. 1986).

Successful control of *S. exigua* is challenging due to its broad host range, rapid growth rate, its migratory dispersal and its ability to rapidly evolve resistance to pesticides (Fu, et al. 2017; Hu, et al. 2021; Huang, et al. 2021). Moreover, the use of conventional chemical pesticides causes health and environmental issues and is generally less accepted (Wheeler 2002; Omkar 2016). Therefore, there is a pressing need for other, more sustainable, strategies to control *S. exigua* and other *Spodoptera* species. A promising approach includes RNA interference (RNAi)-based insect management (Burand and Hunter 2013; Scott, et al. 2013; Renuka, et al. 2017). One of the major challenges is to find target genes for RNAi to control specific pest species or a range of closely related pest species (Li, et al. 2013; Bi, et al. 2016; Tian, et al. 2019). One way to select potential species-specific candidate genes is by carefully analyzing orthologous relationships of homologous genes in related species. Targeting specific gene(s) of single species using RNAi approaches could be an extremely powerful tool to diminish a specific pest outbreak without harming other (closely-related) arthropod species (Price and Gatehouse 2008; Scott, et al. 2013), which often does occur when applying general insecticides (Schulz 2004). Given the high pest potential of many *Spodoptera* species, lineage-specific genes should be identified that can be targeted during pest outbreaks. However, genomic studies have been focused mainly on *S. frugiperda* (Kakumani, et al. 2014; Gouin, et al. 2017) while other *Spodoptera* species have largely been neglected. To address this gap, we present the *S. exigua* genome assembly and official gene set.

In this study we obtained an RNA-sequencing (RNA-seq) profile across all major life stages of *S. exigua*. We performed in-depth analysis of gene expression patterns during the different developmental stages and identified four *Spodoptera*-specific candidate genes for RNAi-based pest management strategies. Furthermore, we produced a *de novo* assembled genome draft of *S. exigua*, based on one female pupa.

## Results

### Genome annotation and comparison to other Lepidoptera genomes

For the *de novo* assembly of the *S. exigua* genome both Oxford Nanopore long read data and Illumina 2×150nt short read data were used. The TULIP assembly strategy was employed to assemble the genome from the long read data (Jansen, et al. 2017). This was followed by two rounds of error correction with the long read data using Racon (Loman, et al. 2015) and two rounds of Pilon polishing with the Illumina data (Walker, et al. 2014; Goodwin, et al. 2015). The total size of the final polished assembled genome was 419 Mb, which was divided over 946 contigs (largest contig = 4.15Mb) with N50=1.1 Mb (Table 1). To confirm the assembly genome size, a k-mer counting approach was used. After counting the 21 and 27 mers in the Illumina data set, the count tables were analyzed with GenomeScope. The genome size as estimated by k-mer counting was ~370 Mbp, which correlated with the Nanopore assembly size (which is slightly larger). The genome size of *S. exigua* presented here, as well as the GC content (given in %), is comparable to other published *Spodoptera* genomes (Table 1).

**Table 1:**
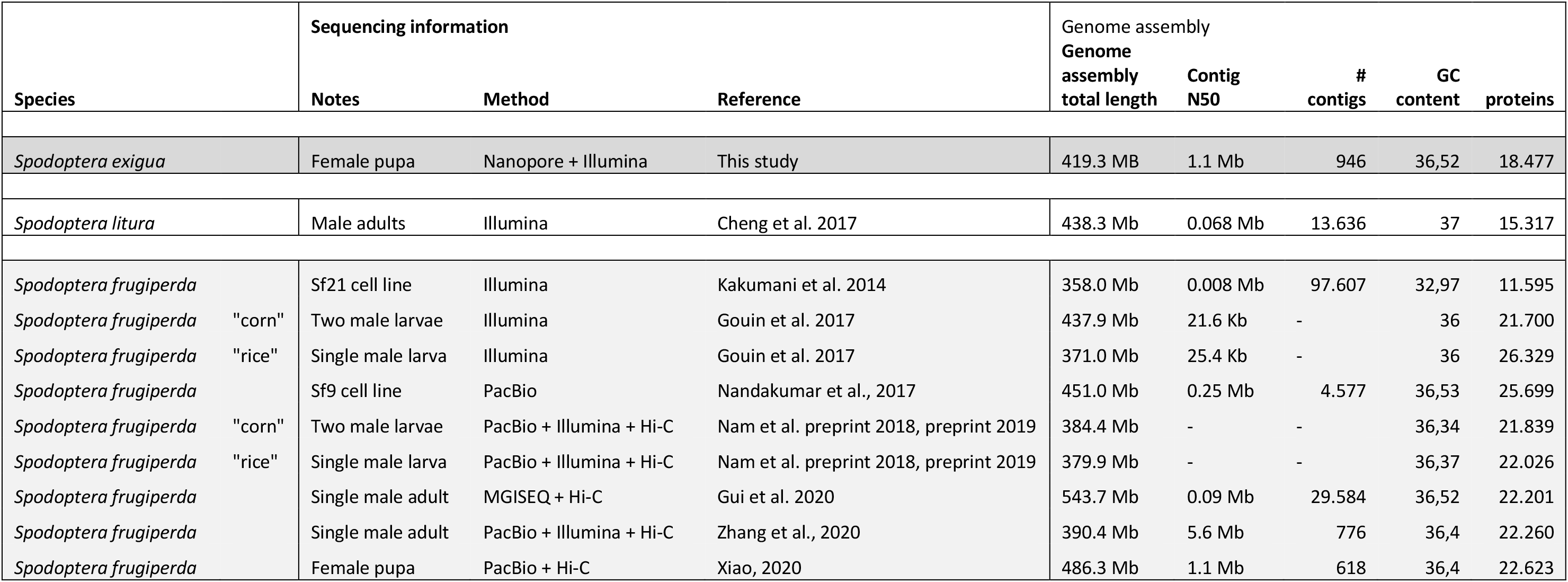
Genome metrics of *Spodoptera exigua* and other published *Spodoptera* genomes.

The Benchmarking Universal Single-Copy Ortholog (BUSCO) (v. 3) assessments indicated that the quality and completeness of our *de novo* assembly was good (complete: 96.8%; fragmented: 1.0%; missing: 2.2%) and comparable to other lepidopteran genomes (Additional file 1: Figure S1). By these quality metrics, the *S. exigua* assembly is comparable to those of fellow lepidopterans, facilitating comparative genomic analyses.

Using our final assembly, an official gene set (OGS) was generated by automatic annotation and transcriptomic RNA-seq data sets of 18 *S. exigua* samples (see below) as supporting evidence. The official gene set, OGS v. 1.1, consists of 18,477 proteins and is provided at the Dryad digital repository.

### Gene expression analyses across the whole life-cycle of Spodoptera exigua

The major developmental stages across the whole life-cycle of *S. exigua*, namely embryonic stage (egg), early 1^st^ instar larva (L1), early 3^rd^ instar larva (L3), pupa, and adult (both sexes: female and male), were sequenced on an Illumina NovaSeq 6000 system at an average of 13.4 million PE2×150nt reads (6.9-22.5 million reads per sample) (Additional file 2: Table S1.3). Based on these reads, we performed differential expression analyses using our *de novo* assembled *S. exigua* genome as a reference.

We first compared gene expression from subsequent different developmental stages and sexes based on pairwise comparisons to determine the dynamic changes in gene expression during development. We used DESeq2 v. 1.18.1 (Love, et al. 2014) to discern significant changes in gene expression throughout the whole life cycle. A striking number of significantly differentially expressed transcripts (n=4,974 transcripts) was detected during early embryonic development (between the egg and the 1^st^ instar larval stage) (Figure 1). Notably, this rapid change in the expression dynamics of *S. exigua* was the largest during the entire life cycle (Figure 1 and Additional file 3: Table S2). In contrast, the smallest change in gene expression was between 1^st^ and 3^rd^ instar larvae (n=1,222 transcripts). A larger change in gene expression was also observed between pupa and adult male (n=3,112 transcripts) compared to pupa to adult female (n=2,061 transcripts), likely due to the fact that female pupae were analyzed. For an overview of relationships between the different life stages based on identified significant changes in gene expression see Additional file 4: Figure S2.

**Figure 1:**
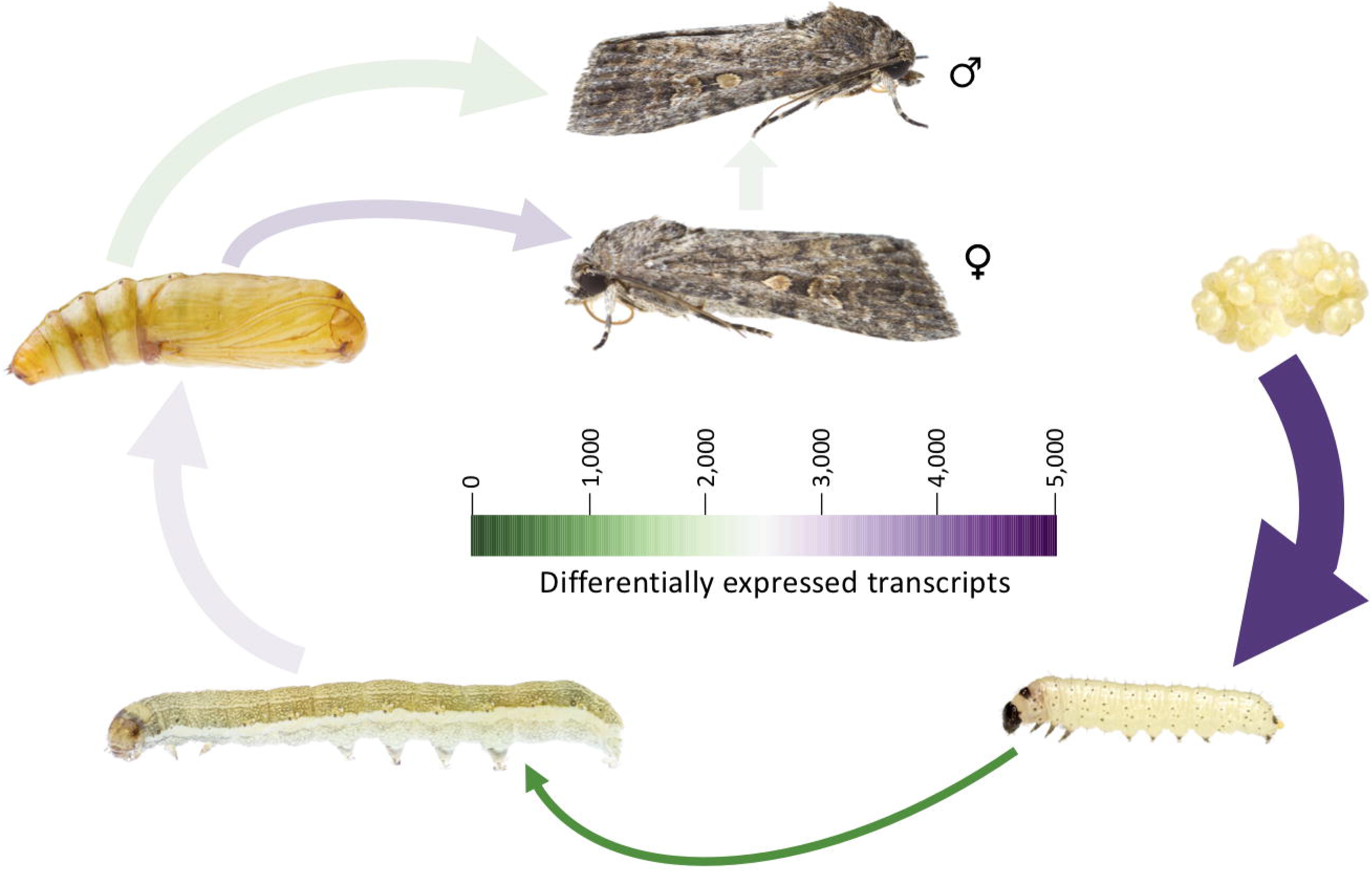
*Spodoptera exigua* life cycle and gene expression profile: The major developmental stages and sexes sequenced for *Spodoptera exigua* are shown, starting from an egg (embryonic stage) and proceeding two larval stages, namely 1^st^ instar and 3^rd^ instar. After the pupal stage, there is the final differentiation into adult male and female. The color of the arrows is proportional to the number of statistically significant differentially expressed genes (FDR = 0.001, minimal fold-change of four), the width an approximation. Note that the size of the developmental stages is not proportional.

We further identified 9,896 transcripts as differentially expressed (DE) across all pairwise comparisons. Hierarchical clustering revealed 14 clusters of DE transcripts with similar expression patterns (Figure 2). Of these, the gene expression of eight clusters could be associated to a single developmental stage or to similar subsequent developmental stages, e.g. one cluster for the larval stage (see also Additional file 5: Figure S3). For these eight clusters, statistically over-represented Gene Ontology (GO) terms were identified using FDR-adjusted *p*-value (< 0.05) and were further summarized to generic GO slim categories (Figure 3).

**Figure 2:**
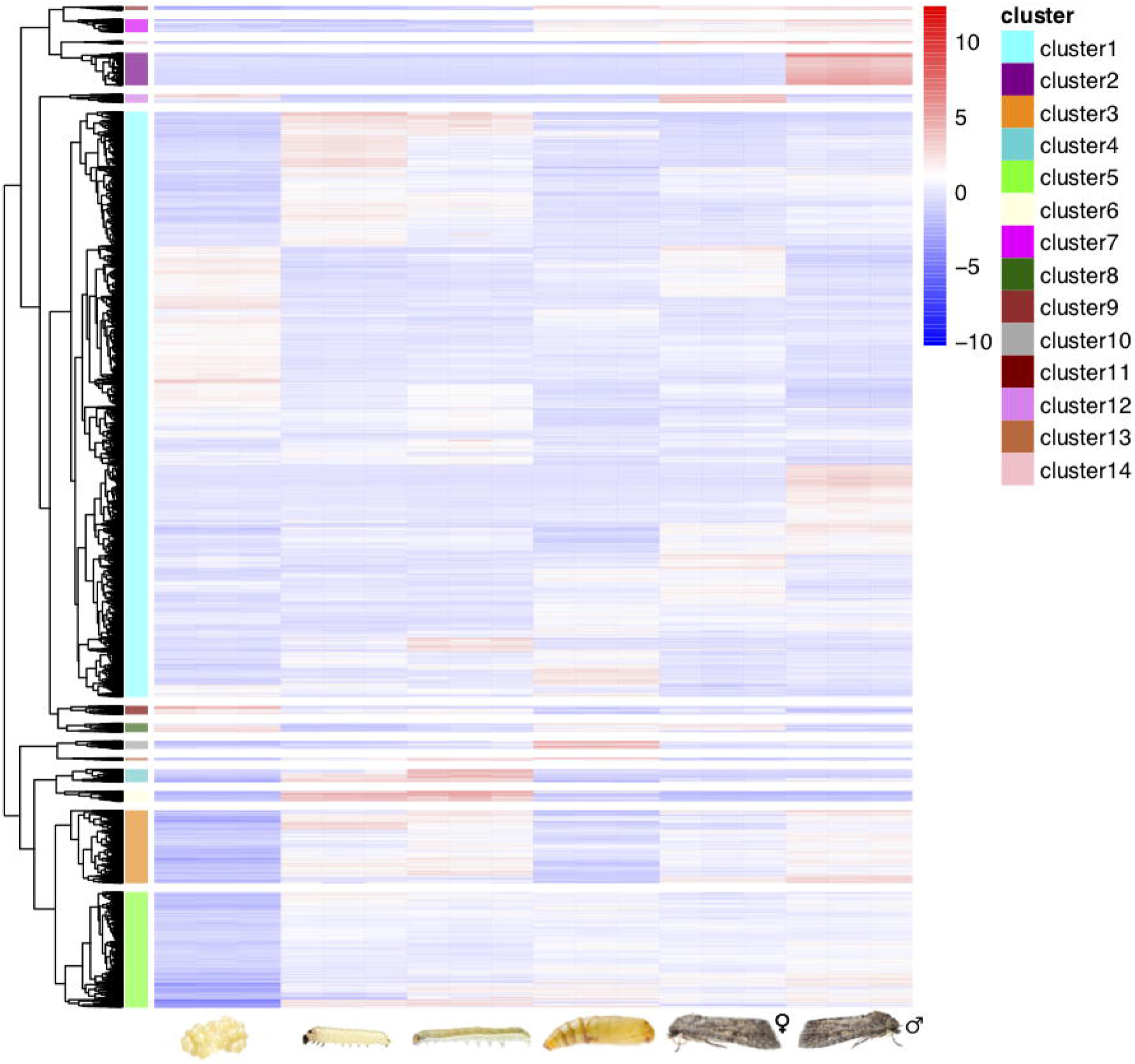
Hierarchical clustering dendrogram of all differentially expressed (DE) genes in the life cycle of Spodoptera exigua: Heatmap shows 9,896 transcripts which have been identified differentially expressed (minimal fold-change of four, FDR ≤ 1e-3) between the six developmental stages / sexes including three replicates each (left to right: embryo, 1^st^ instar larva, 3^rd^ instar larva, pupa, female adult, male adult). Transcripts form 14 distinct clusters using a cut-of at 50% (right dendrogram). The color key of the heatmap indicates low (blue) to high (red) expression values for transcripts.

**Figure 3:**
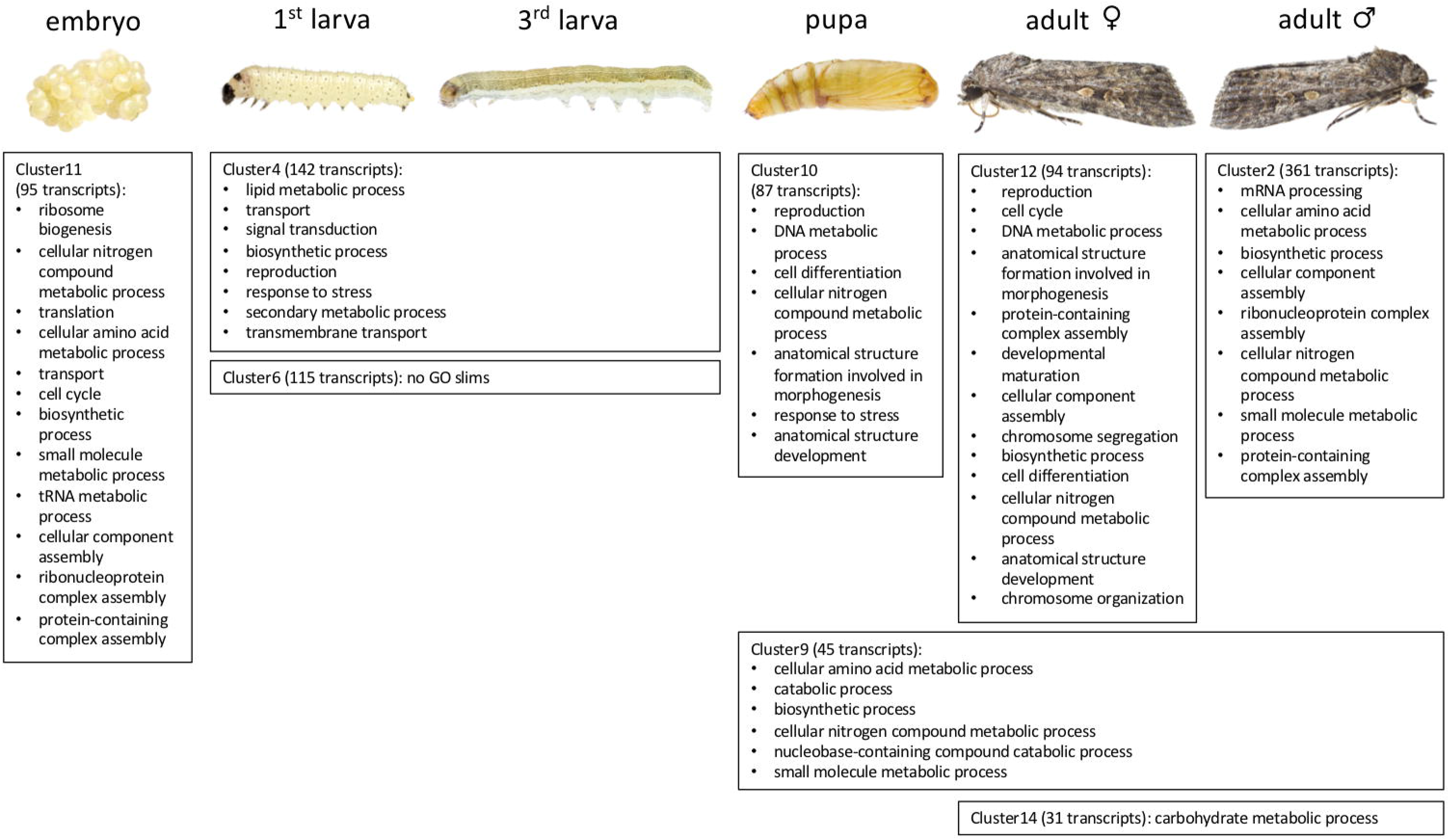
Upregulated GO slims (Biological Process) per development stages: Shown are only the eight clusters of DE transcripts which could be assigned to one developmental stage or sex or to subsequent developmental stages. The cluster number is according to the formed clusters as indicated in Figure 2. The number of transcripts is provided in parentheses as well as the statistically over-represented GO terms (FDR ≤ 0.05) which have been summarized to generic GO slim categories.

For the embryonic stage (cluster 11, Figure 3), there was an enrichment of GO categories associated with ribosome biogenesis (GO:0042254), ribonucleoprotein complex assembly (GO:0022618), tRNA metabolic process (GO:0006399), translation (GO:0006412), and cell cycle (GO:0007049). The enrichment of these categories highlights the rapid succession of cell cycles associated with chromatin replication and initiation of transcription and translation for embryo patterning (Koutsos, et al. 2007). Detailed investigation of DEs gene annotations based on the Arthropoda database (Additional file 6: Table S3 and Additional file 7: Table S4) revealed several known genes important in morphogenesis, for example during the embryonic stage Krüppel-like transcription factors (KLFs) (Kaczynski, et al. 2003; McCulloch and Koenig 2020), specificity proteins (SP) (Kennedy, et al. 2016) and several WD-repeat-containing proteins (Smith 2008).

We did not identify a specific cluster for the 1^st^ larval stage nor for the 3^rd^ larval stage, but rather one cluster including both larval stages (=larval stage cluster, cluster 4, Figure 3). The larval stage was enriched for genes involved in general metabolic processes, such as signal transduction (GO:0007165), biosynthetic processes (GO:0009058), and secondary metabolic processes (GO:0019748). Several genes having a key role in the digestion of plant material and herbivore success were significantly differentially expressed within the larval stage (see Additional file 6: Table S3). These include REPAT genes (Herrero, et al. 2007; Navarro-Cerrillo, et al. 2013), trypsins (Muhlia-Almazán, et al. 2008), cuticle proteins (Celorio-Mancera, et al. 2013; Müller, et al. 2017; Orsucci, et al. 2018; Breeschoten, et al. 2019), and members of prominent detoxification gene families such as cytochrome P450s (P450), carboxyl/cholinesterases (CCE), glutathione S-transferases (GST), and UDP-glycosyltransferases (UGT). The pupal stage varied from the larval stage in that there was significant enrichment in processes associated with cell differentiation (GO:0030154), anatomical structure formation involved in morphogenesis (GO:0048646), and anatomical structure development (GO:0048856).

We further identified several pupal cuticle proteins as significantly differentially expressed within this pupal stage. The female adult stage (cluster 12) was enriched for genes involved in e.g., cell cycle (GO:0007049), chromosome segregation (GO:0007059) and chromosome organization (GO:0051276), anatomical structure development (GO:0048856), and biosynthetic process (GO:0009058) and we identified orthologs of several homeotic genes(-like), such as *Bicaudal C*, *Sex combs reduced* (Scr) and *proboscipedia* (pb). For the male adult stage (cluster 2, Figure 3), there was an enrichment of GO categories associated with e.g. mRNA processing (GO:0006397), cellular amino acid metabolic process (GO:0006520), cellular component assembly (GO:0022607), and biosynthetic process (GO:0009058). For the female and the male adult stage we further identified several sex-specific genes as differentially expressed, such as vitellogenin and vitellogenin receptor in the female (Rotllant, et al. 2017) and testis-specific serine/threonine-protein kinase 2 (Kim, et al. 2019) or ejaculatory bulb-specific protein (Liu, et al. 2020) in the male stage, respectively. One cluster (cluster 14) was specific for both adult sexes but was enriched only for carbohydrate metabolic process (GO:0005975). In contrast, cluster 9 (comprised of the pupa and both adult sexes) was enriched for several GO categories: cellular amino acid metabolic process (GO:0006520), catabolic process (GO:0009056), biosynthetic process (GO:0009058), and cellular nitrogen compound metabolic process (GO:0034641) (see Figure 3 and Additional file 8: Table S5).

### Lepidopteran phylogenomics and detoxification gene content evolution

Phylogenomic analysis correctly placed *S. exigua* within the *Spodoptera* clade and as the sister-group to the clade containing *S. litura* and *S. frugiperda* (Figure 4) (Le Ru, et al. 2018; Kergoat, et al. 2021). In addition, the inferred species relationships within Lepidoptera were in agreement with previous findings (Kawahara, et al. 2019). We further scanned all lepidopteran genomes for gene families associated with detoxification functions. This included: gene families involved in phase I of the detoxification pathway such as cytochrome P450 monooxygenase (P450) and carboxyl- and choline esterase (CCE) (Kant, et al. 2015); gene families involved in phase II, such as UDP-glycosyltransferase (UGT) and glutathione-S-transferase (GST); and the gene family ATP-binding cassette (ABC) involved in phase III (Li, et al. 2007; Heidel-Fischer and Vogel 2015; Kant, et al. 2015). Based on the annotation of the lepidopteran genomes, we searched for expanded detoxification-related genes (Figure 4 and Additional file 9: Table S6). Expansion of major genes families involved in detoxification was mainly visible for *S. frugiperda* (“corn” strain) within the Noctuidae. In the following, we analysed in greater detail several lineage-specific genes.

**Figure 4:**
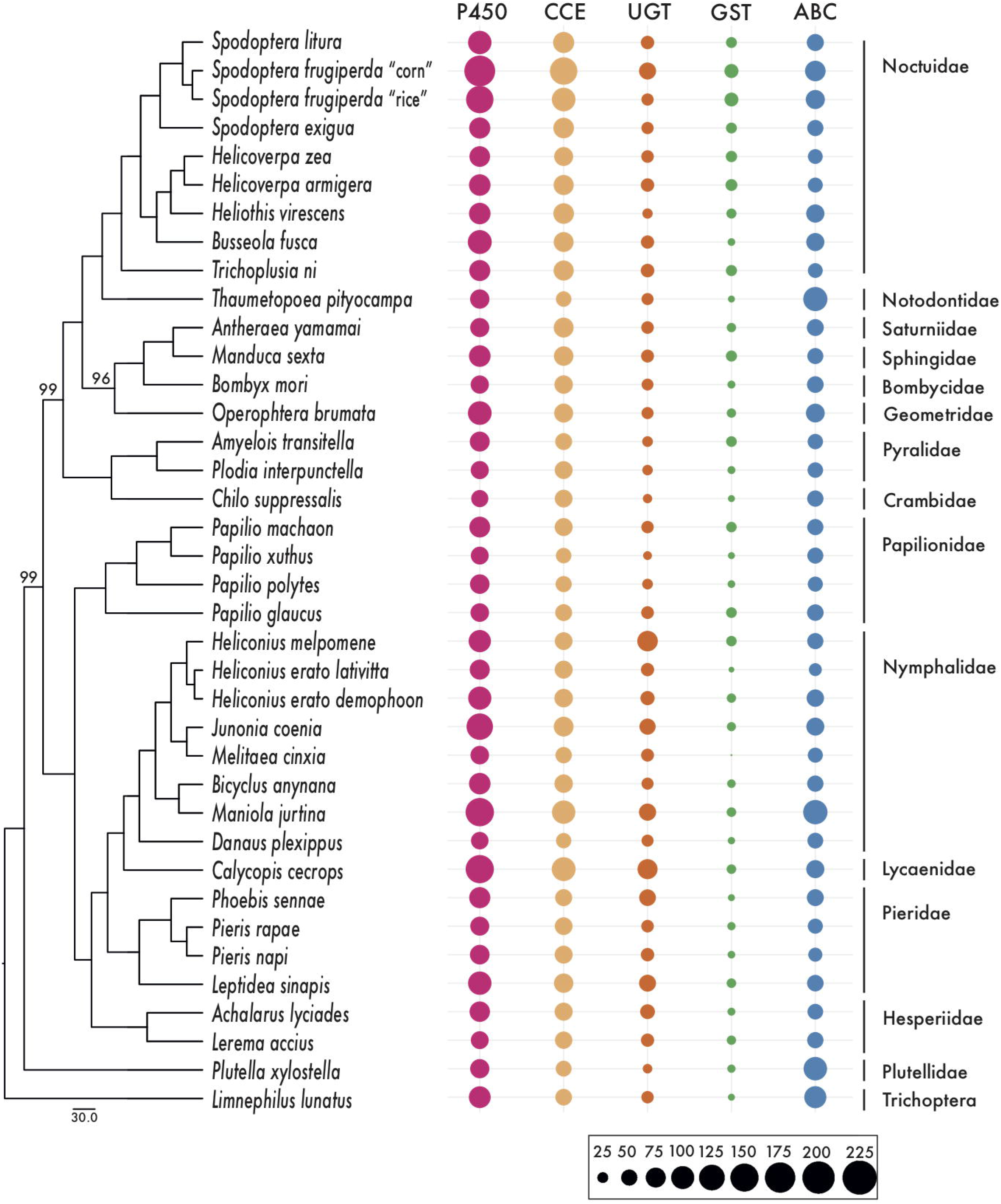
Comparison of Lepidoptera genomes and inferred phylogenetic relationships: Shown is the maximum likelihood phylogeny based on 1,367 single copy BUSCOs (left, all nodes have 100% support unless otherwise noted). Number of detoxification gene members of five main detoxification families, P450 monooxygenases (P450s), carboxyl- and choline esterases (CCEs), UDP-glycosyltransferases (UGTs), glutathione S-transferases (GSTs), ATP-binding cassettes (ABCs), are presented per species in a bubbleplot generated with ggplot2.

### Potential lineage-specific and stage-specific candidate genes as targets for pest-control

We used OrthoFinder v. 2.3.11 (Emms and Kelly 2015) to identify homologous gene sequences in the genomes of eight closely related but diverse lepidopteran species including three *Spodoptera* species, *S. exigua, S. litura,* and *S. frugiperda*. We aimed to identify *Spodoptera*-specific orthogroups, as such lineage-specific genes would be candidates for targeted pest-outbreak management development. We identified in total 119 orthogroups (OGs) containing genes from only the three *Spodoptera* species (Additional file 10: Table S7.1).

Since the larval feeding stage of *Spodoptera* is the most detrimental to crops, we further selected seven OGs for which the *S. exigua* gene representative is differentially expressed in the larval stage cluster (cluster 4). For three of the seven genes, the closest homologs were “uncharacterized” proteins (Additional file 10: Table S7.2). The four remaining genes were annotated as: nuclear complex protein (OG0013351), REPAT46 (OG0014254), trypsin alkaline-c type protein (OG0014208) and mg7 (OG0014260) (Additional file 10: Table S7.2). We confirmed the expression of all seven genes by checking the number of RNA-Seq reads mapped to each assembled transcript based on the results of the transcript abundance estimation with RSEM. The read count in the larval stages (L1 and L3 stages) was higher than in the other stages (Additional file 11: Table S8). Several reads derived from other stages mapped to the protein sequences, but the number of these mapped reads was low (Additional file 11: Table S8).

For the four putative lineage-specific and stage-specific annotated genes, we validated their *Spodoptera*-specificity by constructing gene trees of *Spodoptera* sequences with their most similar sequences identified from other lepidopteran species. For two of the annotated genes (mg7 and REPAT), we constructed two different gene trees that differed in dataset composition. The identification of putative homologs in related species varied per gene as well as the number of included sequences and species for the gene tree analyses (nuclear complex protein (OG0013351): 20 sequences, 3,494 amino acid (aa) positions, REPAT46 (OG0014254) extended dataset containing both αREPAT and βREPAT clusters: 153 sequences, 863 aa positions, reduced dataset containing only the βREPAT cluster: 91 sequences, 717 aa positions, trypsin alkaline-c type protein (OG0014208): 69 sequences, 1,101 aa positions, and mg7 (OG0014260): extended dataset: 27 sequences, 368 aa positions, reduced dataset: 17 sequences, 350 aa positions). All alignment files are provided at the Dryad digital repository. The gene tree of the nuclear pore complex proteins showed that the *Spodoptera*-specific genes form a single cluster, nested within lepidopteran DDB_G0274915-like nuclear pore complex proteins and sister to *Helicoverpa* sequences (Additional file 12: Figure S4). The reduced mg7 dataset comprised sequences from the *Spodoptera*-specific OG in addition to the ortholog group “15970at7088” from OrthoDB. For the extended mg7 dataset we additionally included all ‘mg’ protein sequences according to He, et al. (2012). The ortholog group “15970at7088” included nine single copy genes present in other butterfly species and we found two paralogous copies in *S. litura,* likely due to a specific gene duplication. In order to evaluate whether other paralogs were present in any of the *Spodoptera* gene sets, we blasted the protein sequences against a local blast database of mg7 sequences comprising the sequences from OrthoDB, OG0014260, and He, et al. (2012). In *S. exigua* we identified three paralogs, which according to the GFF file are located (mRNAs) consecutively on the genome: 1268792-1275628, 1276053-1279376, 1280841-1286731. Similarly, in *S. litura* we identified two paralogs and three paralogs in *S. frugiperda*. Running the same blast searches but using the protein sets of *Bombyx mori*, *Helicoverpa armigera*, *Helicoverpa zea*, *Heliothis virescens* and *Trichoplusia ni* all detected a single gene copy with reliable BLAST scores. Both the reduced and the extended mg7 gene trees included all identified *Spodoptera* paralogs. The reduced mg7 gene tree including all paralog *Spodoptera* genes and the single copy homologs from OrthoDB showed that *Spodoptera*-specific OG sequences were clustered together (Additional file 13: Figure S5). This cluster formed a sister clade to all remaining *Spodoptera* paralogs and the *H. armigera* gene. In the extended mg7 gene tree the *Spodoptera*-specific OG sequences did not form a monophyletic clade but did cluster together with the mg7 genes of *Choristoneura fumiferana, H. armigera* and *S. litura* derived from He, et al. (2012) (Additional file 14: Figure S6).

For the REPAT gene analyses we compiled two datasets. Both datasets consisted of sequences derived from the *Spodoptera*-specific OG, the MBF2 ortholog group “16151at7088” from OrthoDB and all protein sequences according to Navarro-Cerrillo, et al. (2013). The reduced dataset only contained protein sequences belonging to the βREPAT class while the extended dataset included both αREPAT and βREPAT classes. In both gene tree analyses, the *Spodoptera*-specific OG sequences clustered together with the annotated REPAT46 gene from *S. exigua* (Additional file 15: Figure S7, Additional file 16: Figure S8). The *Spodoptera*-specific orthogroup is placed in the βREPAT cluster, sensu Navarro-Cerrillo, et al. (2013), where it is placed within group VI (Navarro-Cerrillo, et al. 2013). Further, in total 54 putative REPAT proteins have been identified in the *S. exigua* protein set which were included in both gene tree datasets (Additional file 17: Table S9).

The gene tree of the trypsin proteins showed a monophyletic clustering of all Lepidoptera-derived trypsin genes (Additional file 18: Figure S9). In addition, all *Spodoptera* trypsins were clustered within one monophyletic clade, with the *Spodoptera*-specific orthogroup nested within. Trypsins occurred in all Lepidoptera species in large numbers, thus we compared various OrthoFinder runs under different stringency settings (varying the inflation parameter from 1, 1.2, 1.5 (default), 3.1 and 5) to test the degree of “*Spodoptera*-specificity” of this orthogroup. In all five runs, the OG containing the *Spodoptera* trypsin genes was stable (e.g. lineage-specific) and remained unchanged.

## Discussion

Using a combination of Oxford Nanopore long read data and Illumina short read data for the genome sequencing approach, we generated a high-quality genome and transcriptome of the beet armyworm, *Spodoptera exigua*. These resources will be beneficial for future research on *S. exigua* and other noctuid pest species. The developmental gene expression profile of *S. exigua* demonstrated that the transition from embryo to larva is the most dynamic period of the beet armyworm’s transcriptional activity. Within the larval stage the transcriptional activity was highly similar between early (1^st^) and late (3^rd^) instars, making the early larval stage to an ideal stage for pest-control (see below). Within the larval stage, genes involved in secondary metabolic process (GO:0019748) were uniquely expressed during the whole development (Figure 3). In addition, several prominent genes involved in digestion and detoxification, including cytochrome P450s and UDP-glycosyltransferases, and potential target genes for pest control could be identified which are specifically expressed in the larval stage (Additional file 6: Table S3).

The significant enrichment in the pupal stage in processes associated with anatomical structure development reflects the dramatic structural changes of the larva to the adult (Truman and Riddiford 2019). The identified pupal cuticle proteins within the pupal stage have been reported previously by other studies and reflect the morphological changes in wing disc and the larva-to-pupa metamorphosis (Gu, et al. 2013; Ou, et al. 2014).

The gene expression analyses of the developmental transcriptome of *S. exigua* revealed larval stage-specific upregulated genes (cluster 4, Figure 2&3). These identified genes are strong candidates for targeted RNAi of feeding larvae. Targeted RNAi of genes involved in vital functions of the most important larval stage can be an efficient strategy to minimize the detrimental effect of pest species (Xue, et al. 2012). The larva stages of Noctuidae insects are the most damaging to plants. Our homology search revealed seven *Spodoptera*-specific genes with upregulation in the 1^st^ and 3^rd^ instar larval stages, and highest expression levels in the 3^rd^ instar stage (Additional file 11: Table S8). Four of these seven genes were annotated and we confirmed *Spodoptera*-specificity by gene tree analyses.

One putative *Spodoptera*-specific orthogroup consisted of nuclear pore complex proteins. These proteins are involved in the transport of particles through the nuclear envelope (Alber, et al. 2007). Although the gene tree did not follow well established lepidopteran relationships (Kawahara, et al. 2019), e.g. Noctuoidea nested within Papilionoidea (Additional file 12: Figure S4), all identified *Spodoptera* nuclear pore complex proteins clustered together. This is an initial requisite for potential target genes, showing a clear separation of *Spodoptera*- derived sequences to sequences of other species.

We identified mg7 as potential target gene for RNAi. This gene was previously reported to be highly upregulated in all larval stages in the midgut of *S. litura* with an expression peak after larvae have molted into the 6^th^ larval stage (He, et al. 2012). Our results show a similar pattern with an increased expression towards the 3^rd^ instar larva (Additional file 11: Table S8). Expression in the midgut suggests a role in digestion-related processes (He, et al. 2012). The gene tree based on the reduced dataset showed a clustering of *Spodoptera*-specific mg7 genes (Additional file 13: Figure S5). He, et al. (2012) reported several homologs, mg2, mg7, mg9, mg17, in related species which we included in the extended gene tree reconstruction (Additional file 14: Figure S6). The genes derived from the *Spodoptera*-specific orthogroup form a monophyletic group with the mg7 genes of *C. fumiferana, H. armigera* and *S. litura* derived from He, et al. (2012), establishing orthology of Noctuidae and Tortricidae sequences. The spruce budworm, *C. fumiferana* is a notorious conifer-feeding pest restricted to the Nearctic region where it is considered one of the most destructive insect defoliators (Volney and Fleming 2007; Lumley and Sperling 2010). The extended phylogeny identified homologous clusters (although with low support values) of “mg” genes (mg7, mg17 and mg9) in related lepidopteran species. The close-relationship of additional gene family members from other lepidoptera make mg7 a potential candidate for RNAi-based pest-formation control in a wider range of lepidopteran pest species with the caveat that more work is needed to resolve specificity.

Finally, a strong potential target gene for biocontrol are the αREPAT proteins which are involved in various physiological processes and can be induced in response to infections, bacterial toxins and other microbial pathogens within the larval midgut (Herrero, et al. 2007; Navarro-Cerrillo, et al. 2013). Upregulation of REPAT genes has been identified in response to the entomopathogenic *Bacillus thuringiensis* (Herrero, et al. 2007). In *S. frugiperda,* REPAT genes were associated with defense functions in other tissues than the midgut and found to be likely functionally diverse with roles in cell envelope structure, energy metabolism, transport and binding (Machado, et al. 2016).

REPAT genes are divided in two classes based on conserved domains. Homologous genes of the αREPAT class are identified in closely related *Spodoptera* and *Mamestra* species, while βREPAT class homologs are identified in distantly related species, for example HMG176 in *H. armigera* and MBF2 (Multiprotein bridge factor 2) in *B. mori* (Navarro-Cerrillo, et al. 2013). Our analyses found that REPAT genes (and homologs like MBF2 members) from distantly related species are nested within the βREPAT cluster, while the αREPAT class are exclusive for *Spodoptera* and very closely related species like *Mamestra* spp. (Navarro-Cerrillo, et al. 2013; Zhou, et al. 2016) (Additional file 15: Figure S7, Additional file 16: Figure S8). In contrast to Navarro-Cerrillo, et al. (2013) where αREPAT and βREPAT form sister clades, our tree topology show αREPAT genes to be nested within βREPAT.

Previously, 46 REPAT genes were reported for *S. exigua* (Navarro-Cerrillo, et al. 2013), while we detected 54 REPAT genes in the *S. exigua* genome (Additional file 6: Table S3). The genes of *S. exigua, S. litura* and *S. frugiperda* from the *Spodoptera*-specific orthogroup as identified here cluster together with REPAT46 from *S. exigua* and thus are group VI βREPAT genes (Additional file 15: Figure S7). As shown in Navarro-Cerrillo, et al. (2013) and here (Additional file 15: Figure S7), group VI βREPATs are comprised of *Spodoptera*- and other noctuid-derived genes, like *Helicoverpa* and *Mamestra*. The Noctuidae family is one of the most damaging groups of pests to agriculture, which is recognized by naming of a “pest clade” where species from the genera *Spodoptera*, *Helicoverpa* and *Mamestra* are included (Mitchell, et al. 2006; Regier, et al. 2017). Overall, the results presented here show that REPAT gene members of especially the αREPAT class and the group VI βREPATs are putatively promising candidates for targeted RNAi in notorious pest species belonging to *Spodoptera* and closely related genera in Noctuidae, given their *Spodoptera*- and/or Noctuidae-specificity.

## Conclusions

The genome and developmental transcriptome including all major stages: embryonic, larval, pupal and adult stages of both sexes, of the beet armyworm *Spodoptera exigua* provides a valuable genomic resource for this important pest species. Using a dual sequencing approach including long read and short read data, we were able to provide a genome which is comparable to fellow lepidopterans, strongly supporting the use of these resources in further genome comparisons. Based on the differential gene expression analyses we identified developmental stage-specific (embryonic, larva, pupa or adult) or sex-specific (female, male adult) transcriptional profiles. Of particular interest are the identified genes specifically upregulated in the larval stages because those stages are most detrimental to the host plants. We have further validated these larva-specific genes for their suitability for RNAi-based targeted pest control by comparative genome analyses. RNAi-mediated insect control can be a powerful tool if selected target gene(s) are essential genes in insect tissues to trigger toxic effects. In addition, the target gene(s) should be pest species-specific or specific to a range of closely related pest species and should not harm non-target organisms. In this context, *Spodoptera* lineage-specific target gene(s) are of high interest due to the high number of notorious pest species in this genus causing enormous agricultural damage resulting in economic losses worldwide. Analyzing the homologous relationships of the identified potential target genes and including a broad selection of other insect species allowed us to verify the specificity of four candidate genes for the genus *Spodoptera*. Additional in-depth research may further confirm the clade-specificity of these genes and their potential application in RNAi mediated pest-outbreak management.

## Methods

### Breeding and sample collection

*Spodoptera exigua* specimens originated from a stock rearing of the Laboratory of Virology, Wageningen University & Research, which was initiated in July 2014 using pupae from a large continuous rearing, kindly provided by Andermatt Biocontrol (Switzerland). The rearing was kept on artificial diet at 27°C with 50% relative humidity and a 14:10 h light:dark photoperiod. The artificial diet consisted of water, corn flower, agar, yeast, wheat germ, sorbic acid, methylparaben, ascorbic acid and streptomycin sulphate. Disposable plastic trays covered with paper tissues and a lid were used as rearing containers for groups of maximum 35 larvae (for larger stages). Late 5^th^ instars were transferred to a plastic tray containing vermiculite to facilitate pupation. Pupae were collected and transferred to cylindrical containers lined with paper sheets for egg deposition, with around 45 pupae per cylinder. Adult moths were provided with water. Collected eggs were surface sterilized with formaldehyde vapor to eliminate external microbial contamination.

High molecular weight (HMW) chromosomal DNA was extracted from a female *S. exigua* pupa using the Qiagen Genomic-tip 100/G kit according to the manufacturer’s instructions (Qiagen, Venlo, Netherlands). Quality of the extracted HMW DNA was analyzed on an Agilent 4200 TapeStation System using Genomic DNA ScreenTape (Agilent, Amstelveen, Netherlands). To retrieve samples for RNA sequencing (RNA-Seq), a newly hatched male and female from the continuous rearing were mated in a plastic cup. Offspring of this couple was used for RNA-Seq, six stages were collected: embryonic (eggs), 1^st^ instar larvae, 3^rd^ instar larvae, pupae, male adults, female adults, with three replicates (individuals) per stage except for the embryonic stage were three clusters of each ~100 eggs were taken. To obtain the samples, eggs were harvested, and larvae were reared as above. For the embryonic stage, egg clusters (laid within 21 hours) were cut out of paper, transferred to Eppendorf tubes, snap frozen in liquid nitrogen and transferred to −80 °C until shipment on dry ice to Future Genomics Technologies for further RNA extraction and sequencing. Synchronized newly hatched (non-fed) 1^st^ instar larvae, early 3^rd^ instar larvae, second day pupae and newly emerged (non-mated) female and male adults were collected. Individuals were transferred to Eppendorf tubes and snap frozen as before.

For an overview of all samples please refer to Additional file 2: Table S1. Please refer also to Figure 1 for an overview of the developmental stages.

### Sequencing and assembly of the S. exigua genome

A dual sequencing approach was used for *de novo* assembly of the *S. exigua* genome sequence. In total ~100 Gb of raw Nanopore long read data (Oxford Nanopore Technologies, Oxford, UK) and ~73 Gb of raw Illumina 2×150nt short read data were generated. Long sequence read data was generated using the Oxford Nanopore Technologies platform. Prior to library preparation, HMW DNA was sheared to ~12.5 kb fragments using Covaris gTUBE (Covaris Inc., Woburn, MA, USA). Quality was checked on the Agilent TapeStation. Library preparation was done with the SQK-LSK109 1D ligation kit from Oxford Nanopore Technologies (ONT). Samples were sequenced using one run on an ONT MinION R9.4.1 flowcell and one run on an ONT PromethION R9.4.1 flowcell, respectively. Basecalling was done with Guppy v2.2.2 (ONT MinION) and v1.6.0 (ONT PromethION), respectively. Basecalled reads were used for further processing and assembly.

In addition to long sequence read data, short read data were generated using the Illumina NovaSeq 6000 system. Library preparation was done with the Nextera DNA Flex Library Prep Kit following manufacturers’ protocol (Illumina Inc. San Diego, CA, USA) and quality was checked using the Agilent Bioanalyzer 2100 High Sensitivity DNA Kit (Agilent, Amstelveen, Netherlands). The genomic paired-end (PE) library was sequenced with a read length of 2×150 nucleotides. Image analysis and basecalling was done by the Illumina pipeline. Please refer to Additional file 19: Table S10 for an overview of the DNA sequencing approach. All raw reads from the Illumina, MinION and PromethION sequencing runs were submitted to the NCBI SRA database under accession number PRJNA623582 under sample number SAMN14550570. To assemble the *S. exigua* genome sequence, only long sequence read data were used. First, all reads with a quality score lower than qv=7 were removed from the long sequence read data set. Then, the SEA program (Future Genomics Technologies BV, Leiden, The Netherlands) (Jansen, et al. 2017) was used to prepare seed sequences from the longest reads. In total ~30x estimated coverage of the longest reads were then aligned to these seeds. Reads, alignments and seed files were used to run Tulip v. 1.0.0 (Future Genomics Technologies BV, Leiden, The Netherlands) (Jansen, et al. 2017) to obtain an assembly. The assembly results were used to further optimize the assembly parameters. After this optimization, the total size of the assembled genome was 419 Mb, which was divided over 946 contigs (largest contig = 4.08Mb) with a contig N50 of 1.10 Mb. To further optimize the genome assembly, Racon (Loman, et al. 2015) was used (two rounds) to correct mistakes in the assembly and then two rounds of Pilon polishing (Walker, et al. 2014; Goodwin, et al. 2015) were used to polish the assembly based on the genomic Illumina reads and to reach a high accuracy of the *de novo* assembly that was the basis for genome annotation. The final genome assembly was submitted to the NCBI GenBank database and is available under accession JACEFF000000000, version JACEFF010000000 is used in this study. As a quality check a BUSCO v. 3.0.2 analysis was done on the polished *de novo* assembly using the “insecta_odb9” dataset.

### Sequencing the developmental transcriptome of Spodoptera exigua

Following the Illumina Truseq stranded mRNA library prep protocol (150-750 bp inserts), we prepared 18 different indexed RNA-Seq libraries representing the different developmental stages (namely embryonic stage, early 1^st^ instar larva, early 3^rd^ instar larva, pupa, adult (female and male), and including three biological replicates per stage/sex (Additional file 2: Table S1.1). Libraries were sequenced on an Illumina NovaSeq 6000 system at an average of 13.4 million PE2×150nt reads (6.9-22.5 million reads) per sample at Future Genomics Technologies BV, Leiden, The Netherlands. For an overview of the number of raw reads per samples please refer to Additional file 2: Table S1.3. The sequencing reads were quality checked using FastQC v. 0.10.1 (Andrews 2010). Adapter sequences were removed and quality-filtered using Trimmomatic v. 0.36 (Bolger, et al. 2014), with parameters set: TruSeq3-PE-2.fa:2:30:10, LEADING: 5, TRAILING: 5, SLIDINGWINDOW:4:20, and removing all reads of less than 36 bp in length. All raw reads from the Illumina RNA sequencing approach were submitted to the NCBI SRA database under accession number PRJNA623582.

### Annotation of the Spodoptera exigua genome sequence

The assembled and polished genome was annotated using the maker3 pipeline (maker-3.01.02-beta). As a first step in this analysis a repeat library was constructed with RepeatModeler (RepeatModeler-open-1.0.11; -database Spodoptera_exigua). This species-specific library was used in addition to the RepeatMasker library (Lepidoptera). For gene prediction, Augustus v. 3.3.2 was used which used the model from heliconius_melpomene1 to find genes. As additional evidence for gene models the protein sequences for the family of the Noctuidae were extracted from UniProt (accessed March 7, 2019). Also the RNA-Seq data sets of our 18 *S. exigua* samples were used as supporting evidence. This data set was first assembled using the De Bruijn graph-based *de novo* assembler implemented in the CLC Genomics Workbench version 4.4.1 (CLC bio, Aarhus, Denmark). The available *S. exigua* mRNA nucleotide data from NCBI Genbank (accessed March 7, 2019) was added to this data. After running the pipeline, maker3 annotated a total of 18,477 transcripts. Gene annotations, predicted mRNA and proteins, and assemblies for gene annotations are also provided at the Dryad digital repository.

*Spodoptera exigua* proteins from the official gene set OGS v. 1.1 were further annotated using InterProScan (v. 5.36-75) with several approaches including Go Term annotation (Jones, et al. 2014). Of the 18,477 transcripts, 16,718 transcripts retrieved annotations (Additional file 20: Table S11). Furthermore, the transcript OGS was used in a local BLASTX search v. 2.6.0 (Camacho, et al. 2009) (max_hsps 1, best_hit_overhang 0.1 and E-value cutoff ≤1e-3) against a locally constructed database of all Arthropoda protein sequences downloaded from the NCBI protein database (accessed, 31/01/2019). The translated proteins were additionally used in a BLASTP search v. 2.6.0 (Camacho, et al. 2009) against the same Arthropoda database and parameters (Additional file 6: Table S3, Additional file 7: Table S4).

### Transcript expression quantification

To estimate transcript expression, reads of all samples from each developmental stage were separately mapped to the newly generated *S. exigua* genome (version JACEFF010000000) using Bowtie2 v. 2.3.4 (Langmead and Salzberg 2012). The isoform and gene abundance estimations were done using RSEM v. 1.3.0 (Li and Dewey 2011). A raw (non-normalized) count matrix was created using the perl script ‘abundance_estimates_to_matrix.pl’ implemented in the Trinity v. 2.5.1 package (Grabherr, et al. 2011). The count matrix was cross-sample normalized using the ‘calcNormFactors’ function in edgeR v.3.20.8 (Robinson, McCarthy, et al. 2010) (R v. 3.4.3) using trimmed mean of M values (TMM) (Robinson and Oshlack 2010). See Additional file 21: Table S12 for the raw counts matrix of isoforms in the samples. The normalized count matrix was further filtered by abundance based on count-per-million values (CPM; to account for library size differences between samples) using edgeR v. 3.20.8 (Robinson, McCarthy, et al. 2010). Only genes with a minimum of 5 counts in at least two of the samples were considered expressed and retained in the dataset, see Additional file 23: Table S13.

To measure the similarity of the samples covering the developmental stages and to verify the biological replicates, we implemented the trinity-provided perl script “PtR”. The PCA plot is generated based on the raw non-normalized isoform count matrix which we centered, CPM normalized, log transformed and filtered using a minimum count of 10 (Additional file 24: Figure S10).

The differential expression analysis was performed using DESeq2 v. 1.18.1 (Love, et al. 2014) as implemented in the Trinity package. Transcripts were considered differentially expressed with a minimal fold-change of four between any of the treatments and a false discovery rate (FDR) of p-value ≤ 1e-3. The CPM and TMM normalized expression values of all differentially expressed transcripts were hierarchically clustered and cut at 50% using the Trinity provided script ‘define_clusters_by_cutting_tree.pl’. This resulted in 14 clusters of differentially expressed transcripts with similar expression patterns that were used in the cluster-specific Gene Ontology (GO) analysis.

See Additional file 25: Table S14 for an overview of cluster membership of all 9,896 DE isoforms and Figure 2 and Additional file 5: Figure S3 for expression patterns.

GO analysis was performed using the GOseq package adjusting for transcript length bias in deep sequencing data (Young, et al. 2010) and using the GO annotation retrieved from the Interpro annotation. See Additional file 26: Table S15 for overview of GO annotations within the clusters. GO terms were further summarized to generic GOSlim categories using the R package GOstats (Falcon and Gentleman 2007). For the identified DE genes, statistically over-represented GO terms in each cluster were identified and further summarized to generic GOSlim categories (Additional file 8: Table S5).

### Phylogenomic analyses and comparative genome analyses

We used BUSCO v. 4.0.5 applying the insecta_odb10 as a reference lineage dataset (Seppey, et al. 2019) and comprising in total 1,367 BUSCOSs, to extract single copy complete BUSCOs on the amino acid (aa) level for *S. exigua* and another 37 lepidopteran genomes (Additional file 27: Table S16).

For the phylogenomic analysis, first, amino acid (aa) sequences of single-copy BUSCO genes were separately aligned using MAFFT v. 7.305 (Katoh and Standley 2013) using the L-INS-i algorithm. For the identification of putative ambiguously aligned or randomized multiple sequence alignment (MSA) sections, we used Aliscore v. 1.2 (Misof and Misof 2009; Kück, et al. 2010) on each MSA with the default sliding window size, the maximal number of pairwise sequence comparisons and a special scoring for gap-rich amino acid data (options −r and −e). After exclusion of the identified putative ambiguously aligned or randomized MSA sections with ALICUT v. 2.3 (Kück, et al. 2010), the final MSAs were concatenated into supermatrices using FASconCAT-G v. 1.02 (Kück and Longo 2014). The resulting dataset comprised 1,367 gene partitions and 687,494 amino acid positions.

Prior to the tree reconstruction, the best scoring amino-acid substitution matrix for each gene partition was selected with ModelFinder as implemented in IQ-TREE v. 1.6.12 (Kalyaanamoorthy, et al. 2017). We restricted the search of the best fitting model to eight amino-acid substitution matrices appropriate for nuclear markers: DCMut (Kosiol and Goldman 2005), JTT (Jones, et al. 1992), LG (Le and Gascuel 2008), Poisson, PMB (Veerassamy, et al. 2003), VT (Muller and Vingron 2000), and WAG (Whelan and Goldman 2001). We additionally included the protein mixture model LG4X (Le, et al. 2012), which accounts for FreeRate heterogeneity. Furthermore, we allowed testing the default rate heterogeneity types (E, I, G, I+G, and FreeRates: R) (Yang 1994; Gu, et al. 1995; Soubrier, et al. 2012), with or without empirical rates (−F, −FU) as well as testing the number of rate categories (−cmin 4 – cmax 15). The best model for each gene partition was selected according to the best second-order or corrected Akaike Information Criterion (AICc) score (Hurvich and Tsai 1989). Dataset and partition scheme including selected models are provided at the Dryad digital repository. Phylogenetic relationships were inferred under the maximum likelihood (ML) optimality criterion as implemented in IQ-TREE v. 1.6.12 (Nguyen, et al. 2015; Chernomor, et al. 2016) using the best scoring amino-acid substitution matrix for each gene partition and the edge-proportional partition model allowing partitions to have different evolutionary rates (option -ssp). We performed 50 independent tree searches (25 searches with a random and 25 with a parsimony start tree). The resulting number of unique tree topologies was assessed with Unique Tree v. 1.9, kindly provided by Thomas Wong and available upon request. Node support was estimated via non-parametric bootstrapping of 100 bootstraps replicates in IQ-TREE and mapped onto the ML tree with the best log-likelihood.

We further scanned all these lepidopteran protein sets for several gene families associated with detoxification function, namely P450 monooxygenases (P450s), carboxyl– and choline esterases (CCEs), UDP-glycosyltransferases (UGTs), glutathione S-transferases (GSTs), ATP-binding cassettes (ABCs). We identified the protein families of all proteins by running InterProScan v. 5.36-75 (-appl Pfam --goterms) (Jones, et al. 2014), additionally we ran a local BLASTP against the UniRef50 database (ftp.uniprot.org/pub/databases/uniprot/uniref/uniref50/uniref50.fasta.gz; release version 31/07/2019, accessed 20/08/2019) using an e-value cutoff of 1e-3. Based on these annotations, genes were selected to belong to any of the gene families of interest if it had a match to one of the Uniref50 cluster terms or Pfam– or InterProScan identifiers (Additional file 28: Table S17). The number of detoxification gene members of the five main detoxification families were plotted for each species in a bubbleplot generated with ggplot2 (Wickham 2016) (Figure 4).

### Comparative analysis of Spodoptera-specific genes

We used OrthoFinder v. 2.3.11 using default settings (Emms and Kelly 2015) to identify homologs within the *Spodoptera* clade. We included the genome protein sequence files from three *Spodoptera* species: *S. exigua* (this study), *S. litura* (direct receival OGSv1 28/09/2019 from authors (Cheng, et al. 2017)) and *S. frugiperda* (ftp://ftp.cngb.org/pub/CNSA/CNP0000513/CNS0099235/CNA0003276/Sf_20190612ynM_v1.pep, accessed 20/09/2019; (Liu, et al. 2019)). In addition, we included five closely related but diverse Lepidoptera species: *Heliothis virescens* (ftp://ftp.ncbi.nlm.nih.gov/genomes/all/GCA/002/382/865/GCA_002382865.1_K63_refined_pacbio/GCA_002382865.1_K63_refined_pacbio_protein.faa.gz, accessed 20/09/2019; (Fritz, et al. 2018)), *Helicoverpa zea* (https://data.csiro.au/collections/#collection/CIcsiro:23812v3, accessed 21/08/2019; (Pearce, et al. 2017)), *Helicoverpa armigera* (https://data.csiro.au/collections/#collection/CIcsiro:23812v3, accessed 21/08/2019; (Pearce, et al. 2017))*, Trichoplusia ni* (ftp://www.tnibase.org/pub/tni/tni_protein_v1.fa.gz, accessed 20/09/2019; (Chen, et al. 2019)), and *Bombyx mori* (http://silkbase.ab.a.u-tokyo.ac.jp/cgi-bin/download.cgi, accessed 20/08/2019; (Consortium 2008)).

We identified 119 orthogroups (OGs) containing sequences only from the three *Spodoptera* species (Additional file 10: Table S7.1). Of these 119 OGs, only seven OGs were differentially expressed in the larval stage (cluster 4, Additional file 10: Table S7.2). Of these seven OGs, three OGs were “uncharacterized” protein and four OGS were annotated as: nuclear complex protein (OG0013351), REPAT46 (OG0014254), trypsin alkaline-c type protein (OG0014208), and mg7 (OG0014260) (Additional file 10: Table S7) for which we performed gene tree analyses. For the gene tree analyses, we extended our dataset based on the original OrthoFinder run by including similar sequences from related species to additionally verify the lineage-specificity of these genes. Using the identified *S. exigua* sequences within the lineage-specific OGs as queries, we searched for close homologs using BLASTX (Bravo, et al. 2019) against the NCBI protein database online (Sayers, et al. 2020). Thus, the resulting datasets used to construct gene trees were compiled with some differences. The gene tree of nuclear pore complex proteins was composed of *Spodoptera* orthogroup sequences and all Lepidoptera nuclear complex DDB_G0274915 proteins from the NCBI-nr database (accessed 2/10/2020, keyword “DDB_G0274915”). The initial BLAST identifications of *Spodoptera*-specific orthogroup sequences showed high similarity with DDB_G0274915-like nuclear pore complex proteins. For the remaining three datasets, we additionally included clusters of homologous genes from OrthoDB v. 10 (Kriventseva, et al. 2019). For the REPAT protein dataset, we added the ortholog cluster (“16151at7088”) consisting of “MBF2” orthologs. MBF2 proteins are described to be homologs of REPAT genes in other Lepidoptera species, and have been therefore included (Navarro-Cerrillo, et al. 2013). The REPAT protein gene tree dataset included all protein sequences from Navarro-Cerrillo, et al. (2013). For a second REPAT tree, we only analyzed sequences from the βREPAT class (Navarro-Cerrillo, et al. 2013). For both, the trypsin and mg7 gene tree datasets we included clusters of homologous genes from OrthoDB v. 10 based on the linked cluster to our closest BLAST hit via the online NCBI protein database. For the trypsin gene tree dataset, we added the ortholog cluster “118933at50557” consisting of “serine protease” orthologs. These homologous sequences were selected because the *S. litura* sequence (‘SWUSl0076430’) from the *Spodoptera*-specific orthogroup formed a member of this group. All insect orthologs were included. Finally, the mg7 gene tree dataset included the ortholog group “15970at7088” from OrthoDB v. 10 (accessed 15/09/2020), because the *S. litura* sequence (‘SWUSl0113290’) was an ortholog member. For a second tree we included all genes derived from He, et al. (2012), where the expression of mg7 in the midgut of *S. litura* was studied and homologs in related lepidopteran species were analyzed. Finally, we searched for potential paralogs of all target genes in the protein sets of *S. exigua*, *S. litura* and *S. frugiperda* using BLASTP (max_hsps 1, best_hit_overhang 0.1 and E-value cutoff ≤1e-5) with NCBI-BLAST+ v. 2.6.0 (Camacho, et al. 2009) against a local BlastDB of above gene tree datasets of nuclear pore complex, REPAT, trypsin and mg7 proteins.

For all genes, sequences were aligned using MAFFT v. 7.471 with the L-INS-i method and default settings (Katoh and Standley 2013). Gene trees were reconstructed using IQ-TREE v. 1.6.12 (Nguyen, et al. 2015; Chernomor, et al. 2016) using the maximum likelihood method and implementing bootstrap with 100 replications. The preferred model was applied as based on the model selection (Kalyaanamoorthy, et al. 2017). For the nuclear pore complex gene tree, the best-fit model was “WAG+F+G4”, for REPAT including both αREPAT and βREPAT proteins “WAG+F+R4”, for the gene tree consisting only βREPAT proteins “VT+G4”, for the trypsin gene tree “WAG+F+R5” finally for both mg7 based gene trees “LG+G4”.

The gene trees were rooted dependent of included species and gene composition, aiming for earliest branching genes or species, e.g. by selecting earliest branching lineages from Kawahara, et al. (2019). For the nuclear pore complex protein gene tree, *Papilio xuthus* was used for rooting since it branched early within Papilionidae (Kawahara, et al. 2019). For the REPAT gene tree, we used the same approach as Navarro-Cerrillo, et al. (2013), which rooted the tree using the REPAT-like27 and REPAT-like28 cluster. However, for the limited REPAT gene tree only including βREPAT class genes, we rooted using group V of the βREPAT class according to the first group branching off (Navarro-Cerrillo, et al. 2013). The trypsin tree was rooted using the branch giving rise to a Hymenoptera specific cluster. Finally, the mg7 gene trees were rooted using either *Choristoneura fumiferana* (Tortricidae) (mg9 cluster) or, if absent, *Amyelois transitella* (Pyralidae) (Kawahara, et al. 2019).

## Declarations

### Ethics approval and consent to participate

Not applicable

### Consent for publication

Not applicable

### Availability of data and materials

The final genome assembly was submitted to the NCBI GenBank database and is available under the BioProject PRJNA623582, accession JACEFF000000000, version JACEFF010000000 is used in this study. All raw reads from the Illumina, MinION and PromethION sequencing runs and Illumina RNA sequencing run were submitted to the NCBI SRA database under accession number PRJNA623582.

Further genome datasets and other datasets generated during the current study are provided at the Dryad digital repository.

## Author’s contributions

VIDR and SSi initiated the study. VIDR collected samples. HHJ and RPD performed genome and transcriptome sequencing, *de novo* assembly and automated annotation. TB, SSi, MES further optimized the assembly and annotation and performed differential gene expression, comparative genome and gene tree analyses. TB, SSi, MES, VIDR wrote the manuscript. All authors read and approved the final manuscript.

## Acknowledgements

We thank Els Roode and the late Hanke Bloksma for help with the *S. exigua* rearing and sample collection. We also thank Corné van der Linden for providing *S. exigua* photographs. We thank Entocare for their support to this project.

## Supplementary Information

### Additional file 1: Figure S1

BUSCO assessment of *Spodoptera exigua* compared to other Lepidoptera genomes for ortholog presence and copy number. The relationships are according to Figure 4.

### Additional file 2: Table S1

Overview of developmental stages used for the RNA-Seq approach and sequencing results.

Table S1.1: Developmental stages and samples used for the RNA-Seq approach.

Table S1.2: Quality of developmental stages and samples used for the RNA-Seq approach.

Table S.3: Details on RNA-Seq run results.

### Additional file 3: Table S2

Comparison of differential gene expression between *Spodoptera exigua* developmental stages.

### Additional file 4: Figure S2

Clustered heatmap showing Pearson’s correlation for pairwise sample comparisons based on differentially expressed genes.

### Additional file 5: Figure S3

Visualization of expression patterns with the number of genes per cluster.

### Additional file 6: Table S3

BLASTX annotation report of *Spodoptera exigua* proteins from the official gene set OGS v1.1 using local Arthropoda database.

### Additional file 7: Table S4

BLASTP annotation report of *Spodoptera exigua* proteins from the official gene set OGS v1.1 using local Arthropoda database.

### Additional file 8: Table S5

Go-slims and frequency in respective clusters.

### Additional file 9: Table S6

Proteins annotated using InterProScan for the five main detoxification gene families P450 monooxygenases (P450s), carboxyl- and choline esterases (CCEs), UDP-glycosyltransferases (UGTs), glutathione S-transferases (GSTs), ATP-binding cassettes (ABCs).

### Additional file 10: Table S7

Table S7.1: List of 119 *Spodoptera*-specific orthogroups (OG) with gene IDs.

Table S7.2: Within the 119 OGs, seven *Spodoptera exigua* genes were differentially expressed and clustered in expression cluster 4.

### Additional file 11: Table S8

Read counts of all samples for the seven selected genes based on the ortholog search and specificity for cluster 4.

### Additional file 12: Figure S4

Gene tree of the nuclear pore complex sequences consisting of: the *Spodoptera*-specific orthogroup sequences (OG0013351) and the Lepidoptera sequences from the protein group ‘DDB_G0274915’ as derived from the NCBI protein database. The sequences from the *Spodoptera*-specific orthogroup are marked by the red asterisks, the red frame highlights the *Spodoptera*-specific clade, the red arrow indicates the specific *Spodoptera exigua* differentially expressed gene from expression cluster 4. All sequences derived from the *S.* exigua genome (this study) are underlined. Sequences derived from the protein group ‘DDB_G0274915’ are given their full species names and gene ID.

### Additional file 13: Figure S5

Gene tree of the mg7 sequences (reduced dataset) consisting of: the *Spodoptera*-specific orthogroup sequences (OG0014260) and sequences from OrthoDB cluster ‘15970at7088’. The sequences from the *Spodoptera*-specific orthogroup are marked by the red asterisks, the red frame highlights the *Spodoptera*-specific clade, the red arrow indicates the specific *Spodoptera exigua* differentially expressed gene from expression cluster 4. All sequences derived from the *Spodoptera* exigua genome (this study) are underlined. Sequences derived from the OrthoDB cluster are given their full species names.

### Additional file 14: Figure S6

Gene tree of the mg7 sequences (extended dataset) consisting of: the *Spodoptera*-specific orthogroup sequences (OG0014260), sequences from OrthoDB cluster ‘15970at7088’ and all ‘mg’ gene sequences from He et al. (2012). The sequences from the *Spodoptera*-specific orthogroup are marked by the red asterisks, the red frame highlights the clade, the red arrow indicates the specific *Spodoptera exigua* differentially expressed gene from expression cluster 4. All sequences derived from the *Spodoptera exigua* genome (this study) are underlined. Sequences derived from the OrthoDB cluster are given their full species names only. Genes from He et al. (2012) are given their species names, followed by their gene ID.

### Additional file 15: Figure S7

Gene tree of βREPAT sequences (reduced dataset) consisting of the *Spodoptera*-specific orthogroup (OG0014254), sequences from OrthoDB cluster ‘16151at7088 – “MBF2”’, and βREPAT gene sequences from Navarro-Cerrillo et al. (2013). The sequences from the *Spodoptera*-specific orthogroup are marked by the red asterisks, the red frame highlights the *Spodoptera*-specific clade, while the red arrow indicates the specific *Spodoptera exigua* differentially expressed gene from expression cluster 4. All sequences derived from the *S. exigua* genome (this study) are underlined. Sequences derived from the OrthoDB cluster are given their full species names and cluster ID “MBF2”. Genes from Navarro-Cerrillo et al. (2013) are given the initials of the species names, followed by the REPAT identifier.

### Additional file 16: Figure S8

Gene tree of the REPAT sequences (extended dataset) consisting of: the *Spodoptera*-specific orthogroup sequences (OG0014254), sequences from OrthoDB cluster ‘16151at7088 – “MBF2”’, and REPAT gene sequences derived from Navarro-Cerrillo et al. (2013). Both βREPAT and αREPAT sequences are included in this gene tree. The sequences from the *Spodoptera*-specific orthogroup are marked by the red asterisks, the red frame highlights the *Spodoptera*-specific clade, the red arrow indicates the specific *Spodoptera exigua* differentially expressed gene from expression cluster 4. All sequences derived from the *S. exigua* genome (this study) are underlined. Sequences derived from the OrthoDB cluster are given their full species names and cluster ID “MBF2”. Genes from Navarro-Cerrillo et al. (2013) are given the initials of the species names, followed by the REPAT identifier.

### Additional file 17: Table S9

BLASTP annotated REPAT genes in the *Spodoptera exigua* protein set using reference REPAT sequences from Navarro-Cerrillo et al. (2013).

### Additional file 18: Figure S9

Gene tree of the trypsin sequences consisting of: the *Spodoptera*-specific orthogroup sequences (OG0014208) and sequences from OrthoDB cluster ‘118933at50557’. The sequences from the *Spodoptera*-specific orthogroup are marked by the red asterisks, the red frame highlights the *Spodoptera*-specific clade, the red arrow indicates the specific *Spodoptera exigua* differentially expressed gene from expression cluster 4. All sequences derived from the *Spodoptera exigua* genome (this study) are underlined. Sequences derived from the OrthoDB cluster are given their full species names and gene ID.

### Additional file 19: Table S10

Overview of DNA-Seq approach and sequencing results.

Table S10.1: Results of the Oxford Nanopore long read sequencing run

Table S10.2: Results of the Illumina short read sequencing run

Table S10.3: Results of the trimming and cleaning steps.

### Additional file 20: Table S11

InterProScan annotation report of *Spodoptera exigua* proteins from the official gene set OGS v. 1.1.

### Additional file 21: Table S12

Raw read counts matrix of isoforms in the different samples.

### Additional file 23: Table S13

CPM, TMM cross-sample normalized and filtered count matrix.

### Additional file 24: Figure S10

PCA plot showing sample relationships based on the filtered, CPM-TMM and log2 transformed centered dataset.

### Additional file 25: Table S14

Genes with included in the 14 clusters.

### Additional file 26: Table S15

Cluster membership with GO annotations

### Additional file 27: Table S16

Overview of Lepidoptera genomes and protein sequence files including source location and accession date used for the phylogenomic analyses.

### Additional file 28: Table S17

Gene family identifiers, using InterProScan identifiers and UniRef cluster terms, used to identify putative gene members from families P450, CCE, UGT, GST and ABC.

